# Noise correlations in the owl optic tectum are differentially impacted by unimodal vs crossmodal competitors

**DOI:** 10.1101/2022.07.12.499738

**Authors:** Douglas J Totten, William M DeBello

**Affiliations:** Department of Neurobiology, Physiology and Behavior, Center for Neuroscience, University of California-Davis, Davis, California 95618

## Abstract

Multisensory integration in mammalian superior colliculus and its avian homolog, the optic tectum, is essential for rapid orienting towards salient stimuli. The benefits of combining across modalities is expected to depend in part on interneuronal noise correlations (NCs) and trial-to-trial response variance (Fano factor; FF), yet there is scant data on these topics. Here we used multielectrode arrays (MEAs) to record simultaneously from cohorts of single units in the deep layers of the owl optic tectum (OTd). Stimuli were presented individually or simultaneously as spatially aligned or non-aligned competitors. NC values varied from -0.35 to 0.94, an unexpectedly large range, and decreased only modestly as receptive fields diverged. Spatially aligned bimodal stimuli summed in a largely additive fashion over a wide range of response magnitudes. Most of the observed variance in bimodal NCs and FFs could be accounted for by an additive rule without accounting for internal noise. For non-aligned stimuli, crossmodal competitors decreased FFs, whereas unimodal competitors did not. Most striking, visual competitors decreased NCs for auditory drivers but increased NCs for visual drivers, indicating that the OTd network is differentially wired to process bimodal competitors. In total, these data provide novel descriptive information regarding the correlational structure of OTd neurons underlying multisensory processing.

**SIGNIFICANCE STATEMENT:** In the owl auditory midbrain and its mammalian homologs, little is known regarding the ensemble response properties during multisensory integration. Using microelectrode arrays we found that noise correlations between pairs of neurons evoked by auditory, visual and bimodal stimuli were large and variable. When two non-aligned (competitor) stimuli were presented simultaneously, response reliability and noise correlations were driven down for crossmodal but not unimodal competitors, which suggests distinct processing strategies. This novel finding predicts that visual competitors drive an increase in localization accuracy for auditory targets.

## INTRODUCTION

The avian optic tectum (OT) and its mammalian homologue the superior colliculus (SC) integrate information across sensory modalities to make a topographic priority map that guides orienting movements (Carello and Krauzlis 2004; Knudsen 2011). Across species, much is known regarding how different sensory inputs are integrated in OT/SC when stimuli are spatially aligned and how they compete with each other when misaligned, with emphasis on average response magnitude (Meredith and Stein 1986a, 1986b; Stein, Stanford, and Rowland 2014) and timecourse (Rowland et al. 2007; Whitchurch and Takahashi 2006; Zahar, Reches, and Gutfreund 2009). Yet the performance of OT/SC may also depend on trial-to-trial variance in responses within single neurons (Fano factor; FF) and the variance shared within populations of neurons (noise correlations; NC) (Abbott and Dayan 1999; Jazayeri and Movshon 2006; Pouget, Dayan, and Zemel 2000; Seung and Sompolinsky 1993). NCs can powerfully impact population readout (Averbeck, Latham, and Pouget 2006; Ecker et al. 2011; Kanitscheider, Coen-Cagli, and Pouget 2015; Zohary, Shadlen, and Newsome 1994), but the detriment or benefit depends on the relationship between NCs and signal correlations, in this case, tuning similarity for auditory and visual spatial information. How FFs and NCs manifest during auditory-visual interactions in the OT/SC is unknown.

In both birds and mammals the superficial layers receive direct retinal input (Berman and Wurtz 2008; Kaas and Lyon 2007; Pettigrew and Konishi 1976), while the intermediate/deep layers (OTd) receive auditory information and send outputs to gaze-control circuitry (Knudsen, Knudsen, and Masino 1993; Wurtz and Albano 1980; Wallace, Meredith, and Stein 1993). Focal activation of discrete sites in the OTd topographic map drive orienting movements to the corresponding regions of space (Apter 1946; du Lac and Knudsen 1990; Robinson 1972) (stimulus localization), while lower levels of activity in SCd (Dorris et al. 2002; Muller, Philiastides, and Newsome 2005) or the forebrain structure that connects to OTd (Winkowski and Knudsen 2006, 2007, 2008) bias perceptual thresholds in an attention-like manner (salience detection). Both functions are expected to depend on the rules of multisensory integration at single neuron and population levels.

Rules for how spatially aligned auditory and visual stimuli combine have been elucidated in cat SC. In that system, weaker unimodal stimuli, especially stimuli close to the response threshold for neurons, tend to sum superadditively, while stimuli comfortably within a neuron’s dynamic range sum additively and powerful stimuli sum subadditively (Ghose and Wallace 2014; Meredith and Stein 1996; Stanford, Quessy, and Stein 2005). Rules for how spatially misaligned (i.e. competitors) auditory and visual stimuli combine have been elucidated in both cat SC (Meredith and Stein 1996) and barn owl OT (Mysore, Asadollahi, and Knudsen 2010; Mysore and Knudsen 2011), leading to the hypothesis that the OTd is capable of flexibly identifying the most salient stimulus, independent of modality, in a winner-take-all fashion (Mysore, Asadollahi, and Knudsen 2011; Mysore and Knudsen 2012, 2013).

To better understand how neuronal populations combine auditory and visual signals we recorded simultaneously from cohorts of single units (range 2 – 17) in the OTd of adult barn owls. Auditory and visual spatial receptive fields were mapped and bimodal stimuli were presented at spatially aligned sites or locations displaced by 20° (competitor stimuli). We found that NCs between both nearby and more distant pairs were large and highly variable, and declined modestly with decreasing tuning overlap (signal correlation). Aligned bimodal stimuli drove responses that were typically well matched to a linear combination of the unimodal components, and a substantial portion of the variance in FFs and NCs could be explained through this linear combination. Consistent with previous reports, bimodal competitors reduced the responses to drivers of either modality. The significant new finding is that crossmodal competitors (auditory driver, visual competitor) decreased NCs whereas unimodal visual competitors increased NCs. Thus, the OTd is differentially wired to process unimodal vs bimodal competitors. Because the neuronal pairs on average exhibited high NC, this selective effect predicts an increase in localization accuracy for bimodal competitors.

## METHODS

### Animals, Surgical Preparation and Neurophysiology

#### Animals

Five untrained adult barn owls of both sexes were used in this study. Animals were group housed in large flight aviaries that afforded a wide range of natural behaviors. All husbandry and experimental methods were approved by the University of California, Davis Institutional Animal Care and Use Committee.

#### Surgical Preparation

Electrophysiological recordings were obtained on multiple sessions. Prior to the first recording a stereotaxic head bolt was surgically attached. Owls were food deprived for 24 hours and anesthetized with isoflurane (1-2.5%) and a mixture of nitrous oxide and oxygen (1:1). Skin and feathers on the top of the skull were removed without damaging the facial ruff, the head bolt was attached to the posterior skull using dental acrylic (Jet Denture Repair Powder, Lang Dental Manufacturing Co.) dissolved in Jet Liquid solvent (Lang Dental Manufacturing Co). The remaining exposed skull was covered in a layer of dental acrylic. Wounds were treated with antibiotic ointment, infused with bupivacaine and sutured to abut the acrylic cap. Owls were hydrated with 0.76% saline solution and ketoprofen (1mg/kg) injected intramuscularly, recovered overnight in lab, and returned to their flight rooms the next day.

On the day of recording, owls were anesthetized, wrapped in a soft cloth and craniotomies were opened above the tectal lobes. Owls were removed from anesthesia, transferred to a sound attenuating chamber (Acoustic Systems) and secured to the stereotax (David Kopf Instruments). Once a cohort of single units was isolated for MEA recording, the mixture of nitrous oxide and oxygen was reapplied for the duration of data acquisition.

#### Neurophysiology

Quartz coated platinum-iridium electrodes (0.5-8.0 MΩ, Thomas Recording) were advanced using a concentric seven-electrode microdrive (Thomas Recording), fitted with a pulled glass pipette to bundle the electrodes. Electrodes were independently positioned for single unit isolation in OTd (layers 11-15). Continuous voltage traces were digitized at 25kHz using a Power1401 (Cambridge Electronic Design) and single units were identified offline on Spike2 (Cambridge Electronic Design) using a combination of template matching and clustering based on PCA and waveform measurements (Fig. 1).

**Figure 1.**
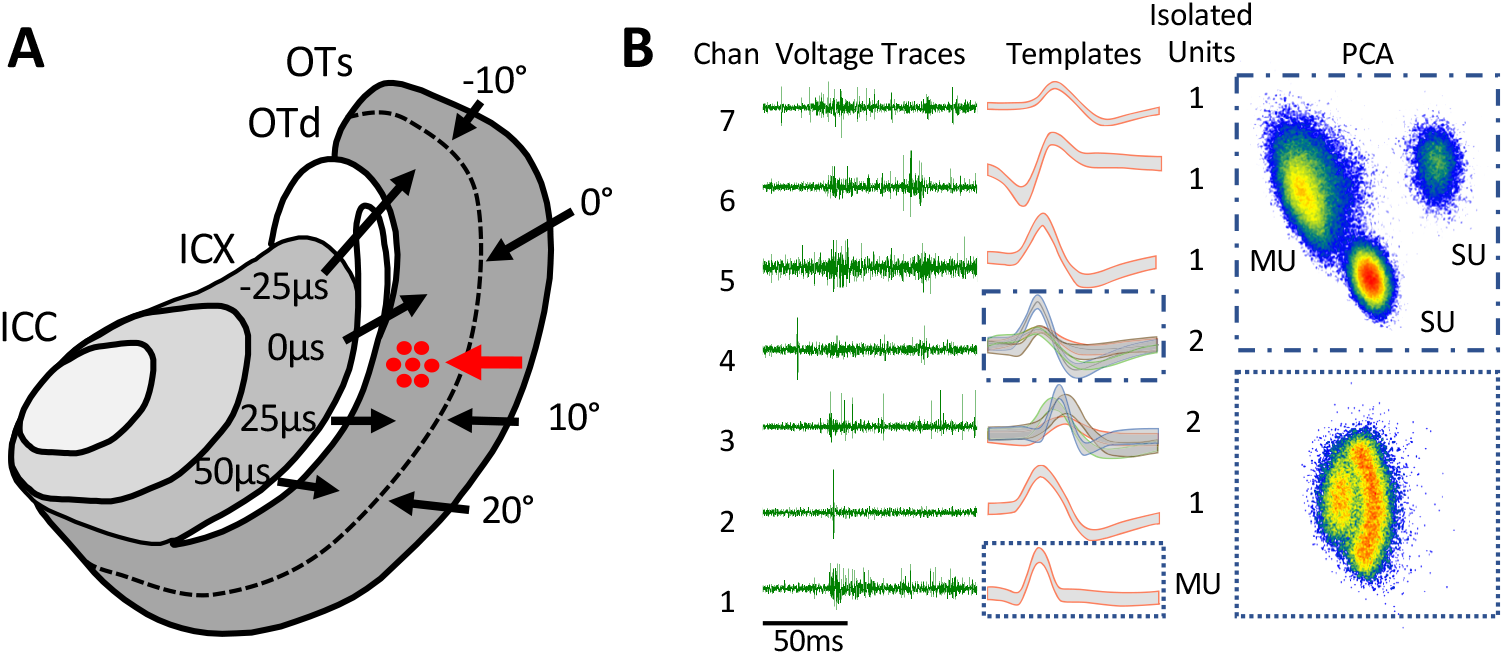
MEA recordings. **A** Diagram of a horizontal section through the midbrain. The map of auditory space is assembled in ICX (external nucleus of the inferior colliculus). Neuronal tuning for ITD is indicated in microseconds. The auditory space map is relayed to OTd where it aligns with a visual space map derived from a retinotopic projection to the superficial layers of OT. Red dots indicate approximate location of MEA trajectories. **B** Voltage traces recorded simultaneously on seven electrodes, with their waveform templates and the number of single units isolated on each channel. Units were isolated using a combination of template matching and principal component analysis (PCA). Waveform heat maps plotted in PCA space for super-threshold events from channels 4 (top) and 1 (bottom) show representative examples of well-isolated single units and inseparable multi-units.

#### Stimuli

Visual stimuli were generated using customized MATLAB software (courtesy of J. Bergan, Harvard University, Cambridge MA) and back-projected (XD221U, Mitsubishi Electric; 60Hz refresh) on a calibrated tangent screen (DA-100, DA-LITE) positioned 58 cm from the owls’ eyes. The owls’ visual axes were aligned by sighting the pectin oculi and all visual stimuli were given in a double-pole coordinate system as previously established (Knudsen 1982). All visual stimuli were static 4° diameter negative contrast filled circle (∼99.4% contrast) presented with a background luminance of 22.5 cd/m^2^.

Auditory stimuli were delivered dichotically through speakers (ED-21913-000, Knowles Electronics) coupled to damping assemblies (BF-1743) positioned ∼5mm away from the tympanic membrane, calibrated to have equalized amplitudes (± 2dB) and aligned frequency-phase portraits. Waveforms were generated using customized MATLAB software (courtesy of J. Bergan, Harvard University, Cambridge MA), interfacing with TDT hardware (RP2 and PA5). All auditory stimuli were broadband frozen noise bursts (2-10khz) with flat amplitude spectra and delivered 20-30dB above auditory threshold. Auditory stimuli were synchronized to projected visual stimuli with a photodiode triggered TTL pulse.

For spatial receptive field (SRF) mapping both visual and auditory stimuli were 100ms in duration with an inter-trial-interval of 1s. Stimuli were presented on a grid with a maximum range of 70° azimuth and 60° elevation of visual space, with spacing of 5°-10° depending on receptive field size and spread, as determined by initial manual mapping. Mapping included from 7-15 repetitions, each consisting of one presentation from each stimulus location, randomly ordered on each repetition. Auditory and visual mapping stimuli were presented in separate blocks.

To measure FFs and NCs, 50 repetitions of each visual, auditory, bimodal and competing stimulus were randomly interleaved. Visual and auditory stimuli were aligned in azimuth using a 2.5μs ITD/degree azimuth transform. Due to the more individually variable relationship between elevation and ILD, visual elevation and ILD were simply set at values that drove responses in as many units as possible given the chosen azimuth/ITD values. A driver location was chosen to drive as much of the cohort as possible based on the auditory and visual RFs measured on each electrode, and the competitor location was placed 20° away along the azimuth dimension. Bimodal stimuli consisted of auditory and visual stimuli at the same location, while competing stimuli consisted of one stimulus of either modality at each location. Each stimulus was 250ms in duration with a 2.5s inter-trial-interval, and all simultaneous stimuli had synchronous onset.

### Data Analysis

#### Response period

The response period was from the time of stimulus onset to the end of the stimulus. This excluded the occasionally large auditory off-response, while still including the majority of the visual response.

#### Spatial tuning

The receptive field weighted average (WA) was calculated as the WA of the peak region. First response rates were baseline-subtracted, then the visual response map was oversampled 10-fold along both spatial dimensions with a cubic spline interpolation. The peak region was defined as the contiguous zone from half-max to maximum interpolated response. The WA was calculated as

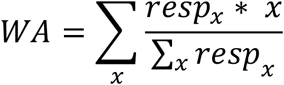

Where *x* corresponds to the stimulus position along one dimension, and *resp*_*x*_ is the response at that position. Because the response maps are two dimensional, the WA was calculated iteratively, by first collapsing across all azimuth values to estimate an elevation value, then taking a cross-section through the response map at the estimated elevation to get an azimuth estimate. The final elevation estimate was taken as the WA of the cross-section at this azimuth estimate, and the final azimuth estimate was taken as the WA of the cross section at the final elevation estimate. These same elevation and azimuth cross-sections were used to calculate the tuning width, which was defined as the full-width of the tuning interpolated response map at half-height.

#### Receptive field dot-product

The receptive field dot-product between units *i* and *j* was calculated using the baseline-subtracted receptive fields with the following equation

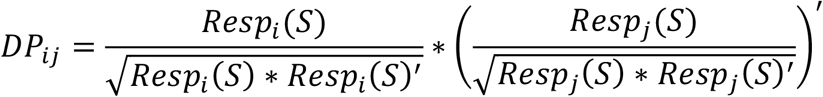

Where *Resp*_*i*_ was the response vector from all spatial locations for unit *i*. Dividing by each unit’s dot-product with itself results in a metric that goes from -1 to 1, with a value of 1 corresponding to a perfect receptive field alignment and -1 corresponding to one receptive field being the inverse of the other (i.e. inhibitory where the other is excitatory and vice-versa).

#### Inclusion criteria

Only units that had significant responses to the static stimulus at the driver location were included in this study. To be included in competition analyses, units also were required to lack a significant response to the stimulus at the competitor location. To determine whether neurons responded to the static stimulus at the driver location, the spike counts during the baseline period before each of the 50 stimulus presentations were used to estimate the expected value of a Poisson-process with those counts. Peristimulus (0-250ms post stimulus) spike trains were then binned into 10ms bins for visual stimuli or 2ms bins for auditory stimuli. The Poisson-expected value was scaled to the bin width and used to run 1000 simulations of a Poisson-generator and find the 95^th^ percentile for expected counts in the bins. Responses were considered significant if they had 3 or more consecutive bins above the 95^th^ percentile.

#### Fano factor and noise correlations

FF and NC were calculated using the single-trial response rates (*R;* not baseline-subtracted) during 50 repetitions of stimuli from a single spatial position. FF was calculated as the variance (*σ*^*2*^*(R)*) in the response rates divided by the mean response rate (*μ(R)*).

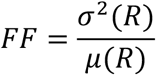

NC was calculated as the Pearson’s correlation coefficient (covariance divided by the product of the standard deviations) between the 50-trial response vectors for each pair of neurons,

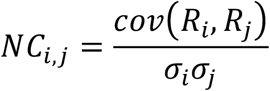

#### Additivity index

Additivity was calculated as previously established (Miller et al., 2015) by taking the difference between the bimodal response and the sum of the unimodal responses, dividing by the sum of the unimodal responses and multiplying by 100 to yield a percent,

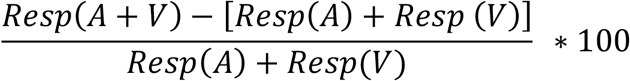

## RESULTS

MEA recordings were targeted to mutually aligned maps of auditory and visual space in the deep layers of OT (OTd). A representative recording is shown in Figure 1. This cohort yielded good waveforms on all seven electrodes from which 8 single units were extracted using offline spike sorting. The complete dataset contained 27 cohorts of 2 – 17 single units (mean = 7.4 units, interquartile range = 3.25 units) recorded from 5 owls (Table 1). Different numbers of units passed inclusion criteria for each analysis described below.

**Table1.** Cohort unit counts. This table includes the owl that each cohort was recorded in, the number of single units that were isolated, the number of those units with significant responses to visual stimuli at the driver location (Vd), auditory stimuli at the driver location (Ad), and units that don’t have significant responses to visual stimuli at the competitor location (∼Vc) or auditory at the competitor location (∼Ac).

| Owl ID | Total Sus | Vd | Ad | Vd+Ad | Vd+~Vc | Vd+~Ac | Ad+~Ac | Ad+~Vc |
| --- | --- | --- | --- | --- | --- | --- | --- | --- |
| 206 | 12 | 3 | 5 | 2 | 3 | 3 | 3 | 5 |
| 206 | 4 | 0 | 2 | 0 | 0 | 0 | 1 | 2 |
| 270 | 5 | 4 | 4 | 3 | 4 | 3 | 3 | 4 |
| 270 | 9 | 4 | 6 | 4 | 4 | 4 | 6 | 5 |
| 206 | 11 | 3 | 6 | 3 | 3 | 3 | 6 | 6 |
| 206 | 7 | 2 | 4 | 2 | 2 | 2 | 2 | 4 |
| 206 | 8 | 4 | 6 | 3 | 4 | 4 | 4 | 4 |
| 281 | 17 | 12 | 13 | 10 | 10 | 6 | 5 | 12 |
| 281 | 8 | 5 | 6 | 3 | 3 | 1 | 2 | 5 |
| 281 | 9 | 6 | 6 | 5 | 4 | 3 | 2 | 4 |
| 281 | 6 | 4 | 6 | 4 | 3 | 2 | 2 | 4 |
| 281 | 11 | 4 | 3 | 2 | 3 | 4 | 3 | 3 |
| 281 | 7 | 5 | NA | NA | 5 | NA | NA | NA |
| 245 | 9 | 4 | NA | NA | 4 | NA | NA | NA |
| 245 | 5 | 4 | NA | NA | 4 | NA | NA | NA |
| 245 | 5 | 3 | NA | NA | 3 | NA | NA | NA |
| 245 | 7 | 3 | NA | NA | 3 | NA | NA | NA |
| 245 | 8 | 6 | NA | NA | 6 | NA | NA | NA |
| 239 | 8 | 1 | NA | NA | 1 | NA | NA | NA |
| 239 | 8 | 3 | NA | NA | 2 | NA | NA | NA |
| 239 | 7 | 0 | NA | NA | 0 | NA | NA | NA |
| 239 | 5 | 2 | NA | NA | 2 | NA | NA | NA |
| 239 | 3 | 2 | NA | NA | 1 | NA | NA | NA |
| 239 | 2 | 1 | NA | NA | 1 | NA | NA | NA |
| 239 | 6 | 2 | NA | NA | 2 | NA | NA | NA |
| 239 | 5 | 0 | NA | NA | 0 | NA | NA | NA |
| 239 | 8 | 2 | NA | NA | 2 | NA | NA | NA |
| 239 | 8 | 5 | NA | NA | 5 | NA | NA | NA |

### Auditory and visual spatial receptive fields

Auditory and visual spatial receptive fields (SRFs) averaged across 26 single units are shown in Figure 2. For this analysis, units were included only if they had strong responses to both auditory and visual stimuli and their receptive fields were well contained within a complete, fine-grain 2D mapping space for both modalities. To facilitate comparison the azimuthal location of the auditory stimulus is displayed in degrees, not ITD, according to the standard transformation of 2.5 μs per degree. Consistent with previous results (Knudsen 1982), auditory SRFs were wider than visual SRFs along the horizontal axis (half-max widths, 14.5° and 9.9° respectively; p < 0.01, n = 26, two-tailed T-test).

**Figure 2.**
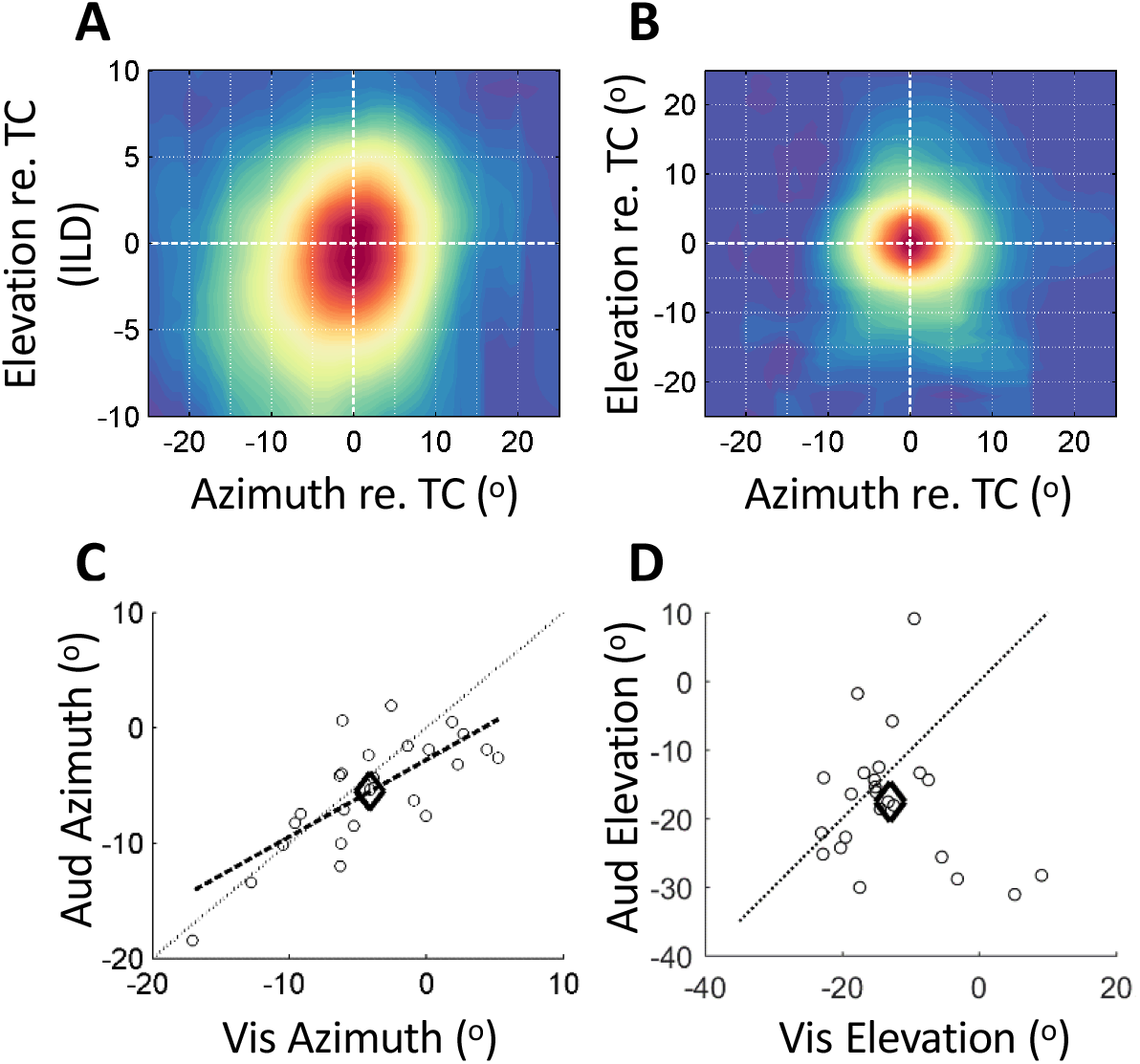
Auditory and visual tuning. **A** Auditory spatial receptive fields were aligned to their tuning centers and averaged (n = 24). ITDs were converted to degrees azimuth (x-axis) but ILDs were kept as the unit of measure for elevation. **B** Same conventions as **A** for visual spatial receptive fields (n = 26). Both azimuth and elevation are in units of degrees. Visual spatial receptive fields were significantly narrower in azimuth than auditory SRFs (Aud = 14.5°, Vis = 9.9°; n = 26, t-test p < .01). **C** Auditory and visual tuning centers were well aligned in azimuth. Small circles indicate single sites (n=26), while the large diamond indicates the mean (Aud = -5.51, Vis = -4.07; t-test p >0.05) and the dashed line indicates the linear fit (rho = 0.76, p < 0.01). **D** Auditory and visual elevation tuning centers were not significantly correlated (rho = -0.24, p > 0.05).

Auditory and visual receptive fields were well aligned in azimuth (Fig. 2C). At the population level, the weighted averages of the auditory and visual spatial receptive fields (SRFs) were not significantly different from each other along the azimuth dimension according to a two-tailed T-test (Fig. 2A), and were well described by a linear relationship. Auditory and visual SRF weighted averages were not correlated in elevation (Fig. 2D). This weak correspondence between ILD and visual elevation likely reflects variability in individual HRTFs, which were not measured.

Bundled electrodes allowed for isolation of single units that typically displayed overlap in their SRFs for both visual and auditory stimuli, as shown in the representative cohort displayed in Figure 3. This allowed targeting of a single spatial location (Fig. 3B and D) from which visual, auditory and bimodal stimuli effectively drive most (rarely all) units in the cohort. Raster plots for the cohort shown in Fig. C, D are shown in Figure 4. Onset latencies to visual stimuli were typically 40-50ms and the responses were often sustained throughout the stimulus. Onset latencies to auditory stimuli were typically 11-14ms and the responses tended to diminish throughout the stimulus. These response profiles are typical of neurons in OTd which inherit their auditory tuning from space-specific neurons (SSNs) in the external nucleus of the inferior colliculus (ICX), and visual tuning via dendritic extensions into the superficial layers of OT which receive a direct retinotopic projection.

**Figure 3.**
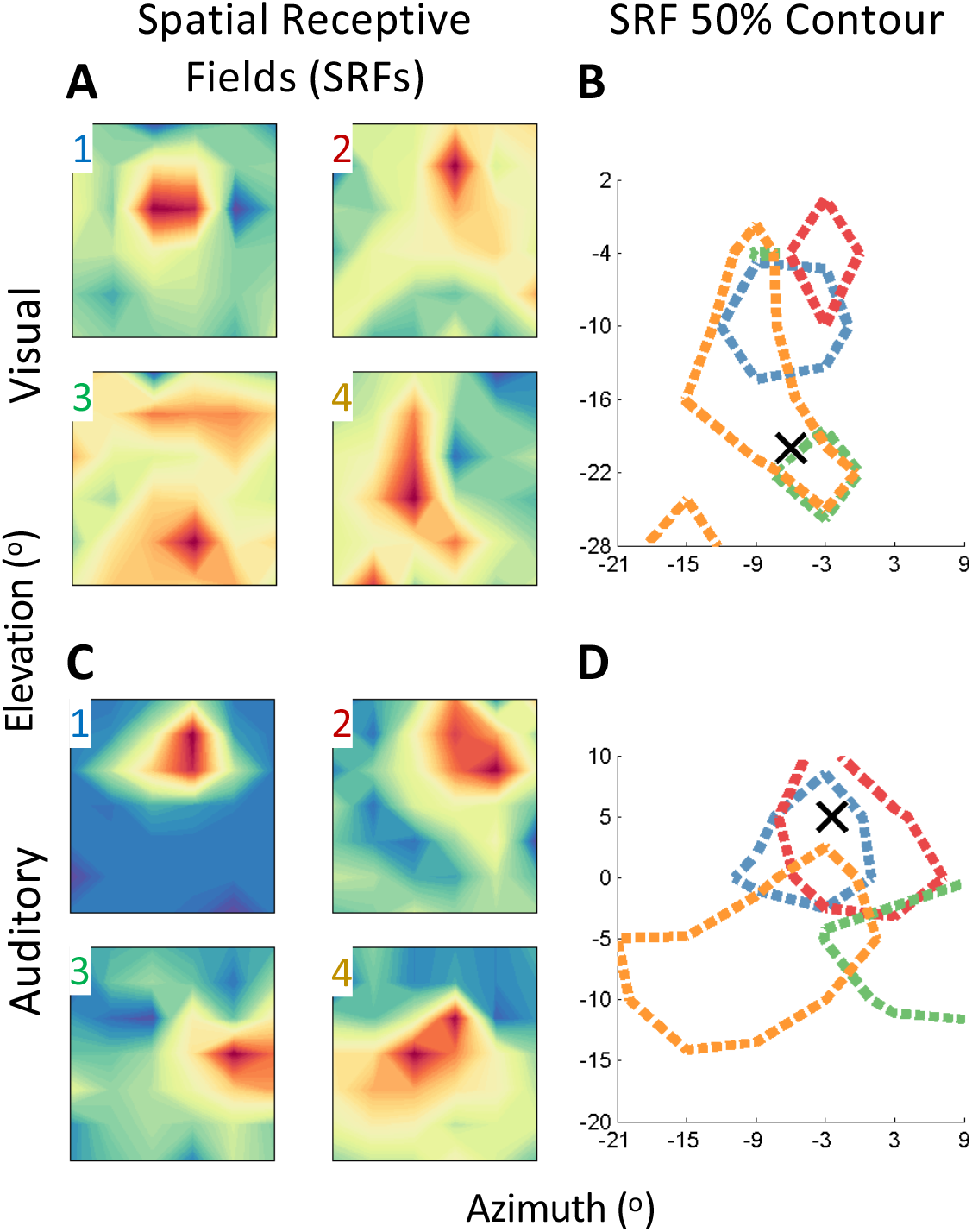
Representative examples of spatial receptive field mapping. **A** Visual SRFs were mapped using negative contrast 4° static dots. **B** For FF and NC assessment a single stimulus location (referred to here as the driver location) was chosen in the overlapping region of the receptive fields. Dotted lines show the 50% maximum response contour, and the X indicates the stimulus location chosen for FF and NC measurements. **C**, Auditory SRFs were mapped using noise bursts 20-30dB above threshold. **D** Similar to **B** except with auditory stimuli.

**Figure 4.**
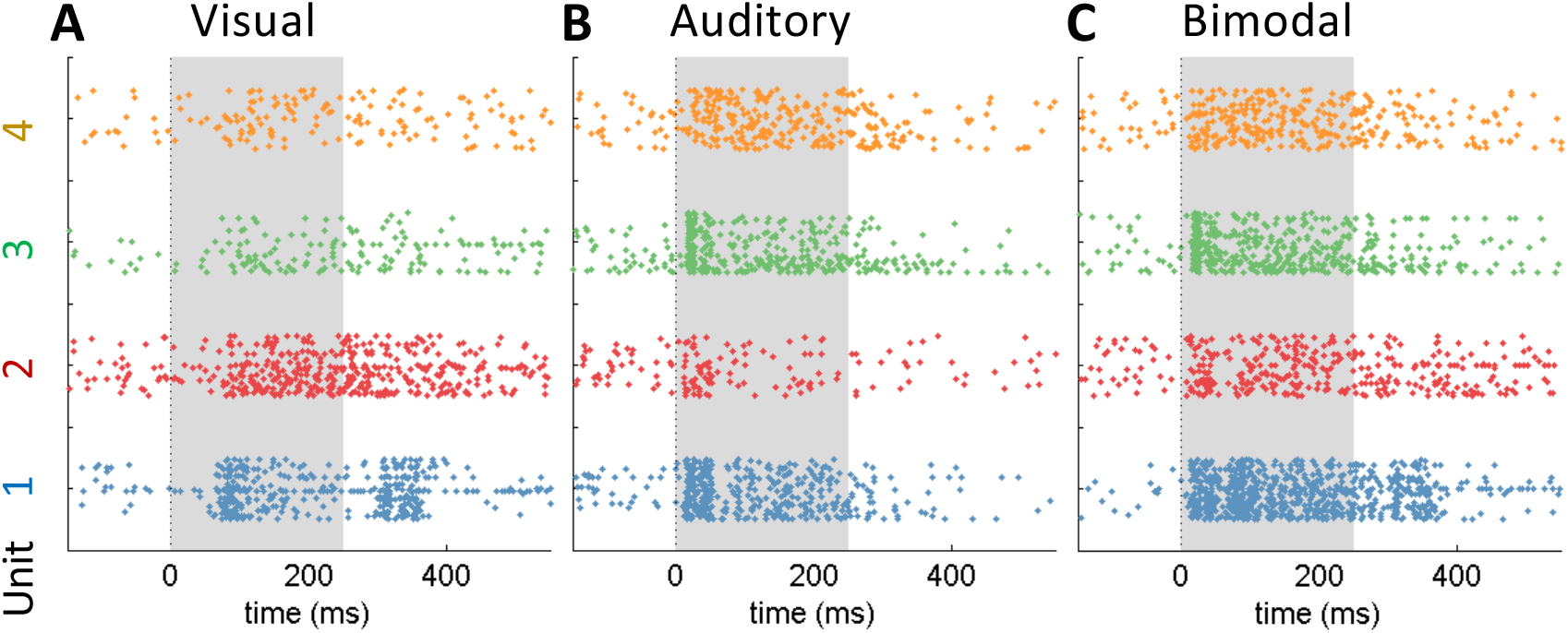
Representative example of response time-course. Raster plots for the responses to **A** (auditory), **B** (visual) and **C** (bimodal) stimuli played at driver location for four representative neurons. Stimulus presentation period is shown as the grey window extending from 0-250 ms.

### Noise correlations and dependence on tuning similarity

NCs evoked by auditory or visual stimulation were measured for 182 and 181 neuronal pairs, respectively. Values ranged from -0.30 to 0.80 (auditory) and -0.22 to 0.93 (visual), a range consistent with a previous report of 14 spontaneously active pairs in owl OT (Netser, Dutta, and Gutfreund 2014). Overall, the mean noise correlation evoked by auditory or visual stimulation was 0.28 and 0.25, respectively. These values are larger than typically observed in mammalian cortex in response to various stimuli (typically < 0.2; see Discussion) and somewhat larger than those observed in ketamine-anesthetized owl OT in response to auditory stimuli (Beckert, Pavao, and Pena 2017). To our knowledge there are no prior reports of noise correlations to auditory and bimodal stimuli in the mammalian homolog structure the superior colliculus.

The magnitude of the NCs between a pair of units is typically related to their tuning similarity (Bair, Zohary, and Newsome 2001; Cohen and Newsome 2008; Jeanne, Sharpee, and Gentner 2013). We used the normalized dot product between the spatial receptive fields for each neuronal pair as a metric for spatial tuning similarity, as it is a model-independent measure for receptive field overlap (Usrey, Reppas, and Reid 1999). Figure 5 shows that the NC values measured from many repetitions at the driver location were related to the SRF dot products for both visual drivers paired with visual SRFs (Pearson’s r = 0.28, p = 0.0001, n=181) and auditory drivers paired with auditory SRFs (Pearson’s r = 0.24, p = 0.0009, n = 182). While this correlation was significant, units with highly overlapping receptive fields could exhibit large, small or even slightly negative noise correlations; units with almost no overlap exhibited nearly as large a range.

**Figure 5.**
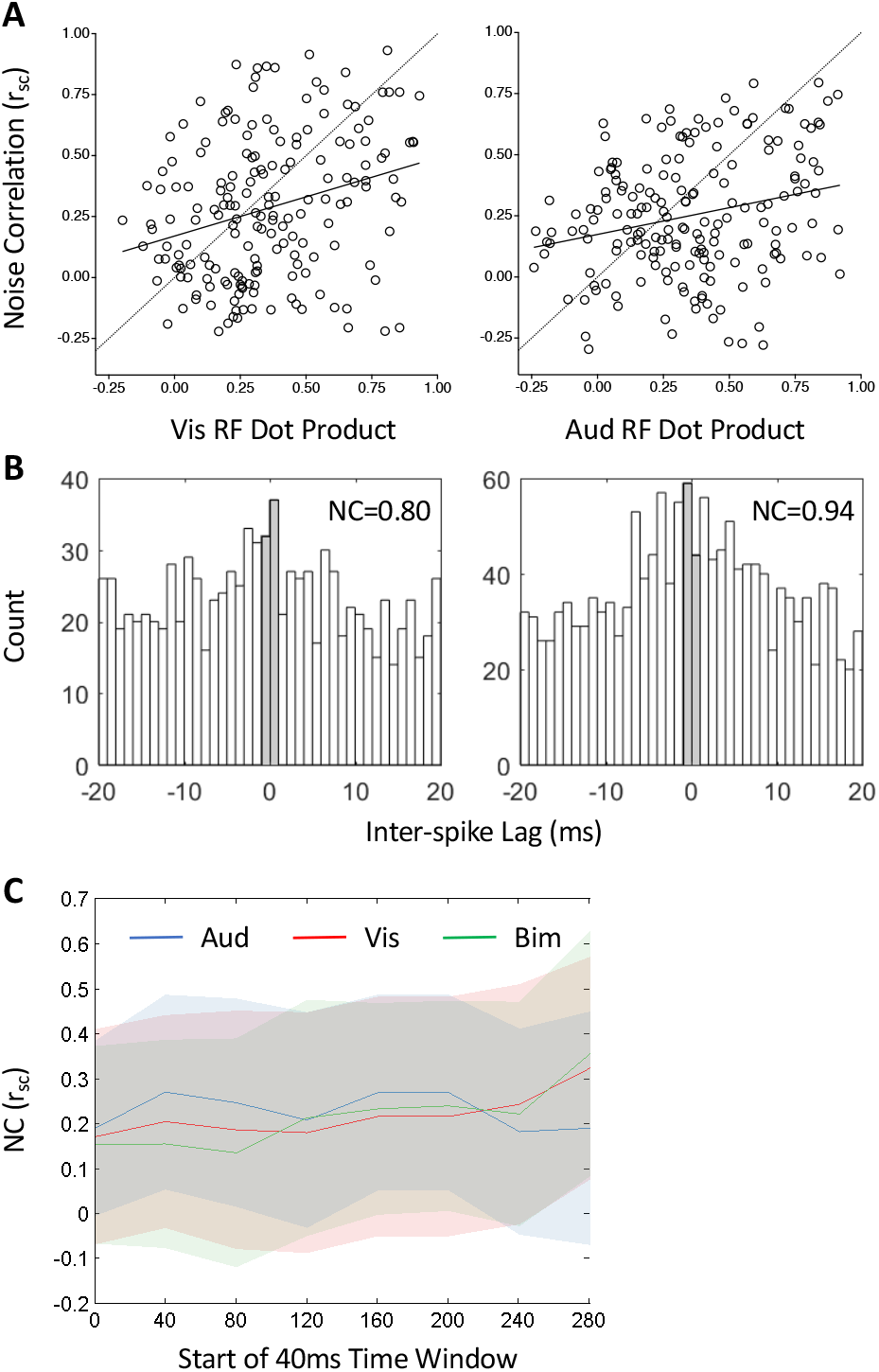
Noise correlations for auditory and visual stimuli. **A** NCs evoked by visual stimulation increased modestly with increasing overlap of receptive fields (Vis RF dot product) (n = 181, rho = 0.28, p < 0.01). The large diamond represents the mean and the dashed line represents the linear fit. NCs evoked by auditory stimulation exhibited a similar trend (n = 182, rho = 0.24, p < 0.01). **B** Cross-correlograms of units recorded on neighboring electrodes, for the two pairs with the highest NC values during the bimodal stimulation condition. Grey bars are bins spanning -1ms:1ms lag difference where correlation should be evident if different electrodes were recording the same unit. **C** Noise correlation time course. Noise correlations were assessed based on spike counts in an 80ms window at 40ms intervals post stimulus onset for auditory (blue) visual (red) and bimodal (green) stimuli.

Large NC values could result from spike sorting errors or from recording the same neuron on multiple electrodes. To address this, we investigated two pairs with the highest NC values in the bimodal stimulation condition (NC = 0.80 and NC = 0.94). In both instances, the units in the pair were isolated on different electrodes and could not result from a spike sorting error. If the same units were detected on multiple electrodes the two units should have large numbers of simultaneous events. The cross-correlograms for the spike trains of these two pairs show that any spikes detected on both channels had a negligible contribution compared to the spikes that were unique to each channel (Figure 5B) i.e. they were different units.

Because auditory and visual responses have distinct onset latencies and temporal response patterns we examined how NCs evolve over time. NCs were calculated in an 80ms window advanced in 40ms intervals, with the start of the window moving from synchronous with stimulus onset to 30ms post stimulus. Figure 5C shows that NCs were stable throughout stimulus presentation. They are lower during this analysis than the others because the short time window leads to lower firing rates which lead to lower correlation values as previously established (Cohen and Kohn 2011). There was a jump in NCs in the last time bin (280ms-360ms; stimulus ends at 250ms) for the visual and bimodal NCs, corresponding to correlations in the off-response. This period is not included in the response-driven NCs presented in all other analyses.

### Linear integration of auditory and visual responses

We first examined the firing rates evoked by auditory, visual and aligned bimodal stimuli. As shown in Figure 6A, auditory stimuli drove larger response rates, on average, than visual stimuli (means = 13.8Hz and 8.2Hz, p = 0.006 with Signed Rank test, n = 40), while bimodal stimuli drove larger response rates than either auditory or visual stimuli presented alone (p < 0.0001 for both with signed rank test). To determine the rules of integration, bimodal responses were compared to the sum of the unimodal responses (Fig. 6*B*). On average, bimodal responses closely conformed to an additive relationship Resp(Aud) + Resp(Vis) = Resp(Aud& Vis)*0.98 + 0.11, Pearson’s r = 0.98). Still, 16/40 units exhibited statistically significant nonlinear response (filled circles in Fig 6B, C and D). The magnitude of the nonlinearity was typically small with a handful of sites exhibiting more than 50% supralinearity. Nonlinear behavior was not predicted by either bimodal strength (Fig. 6C) or unisensory imbalance (Fig. 6D).

**Figure 6.**
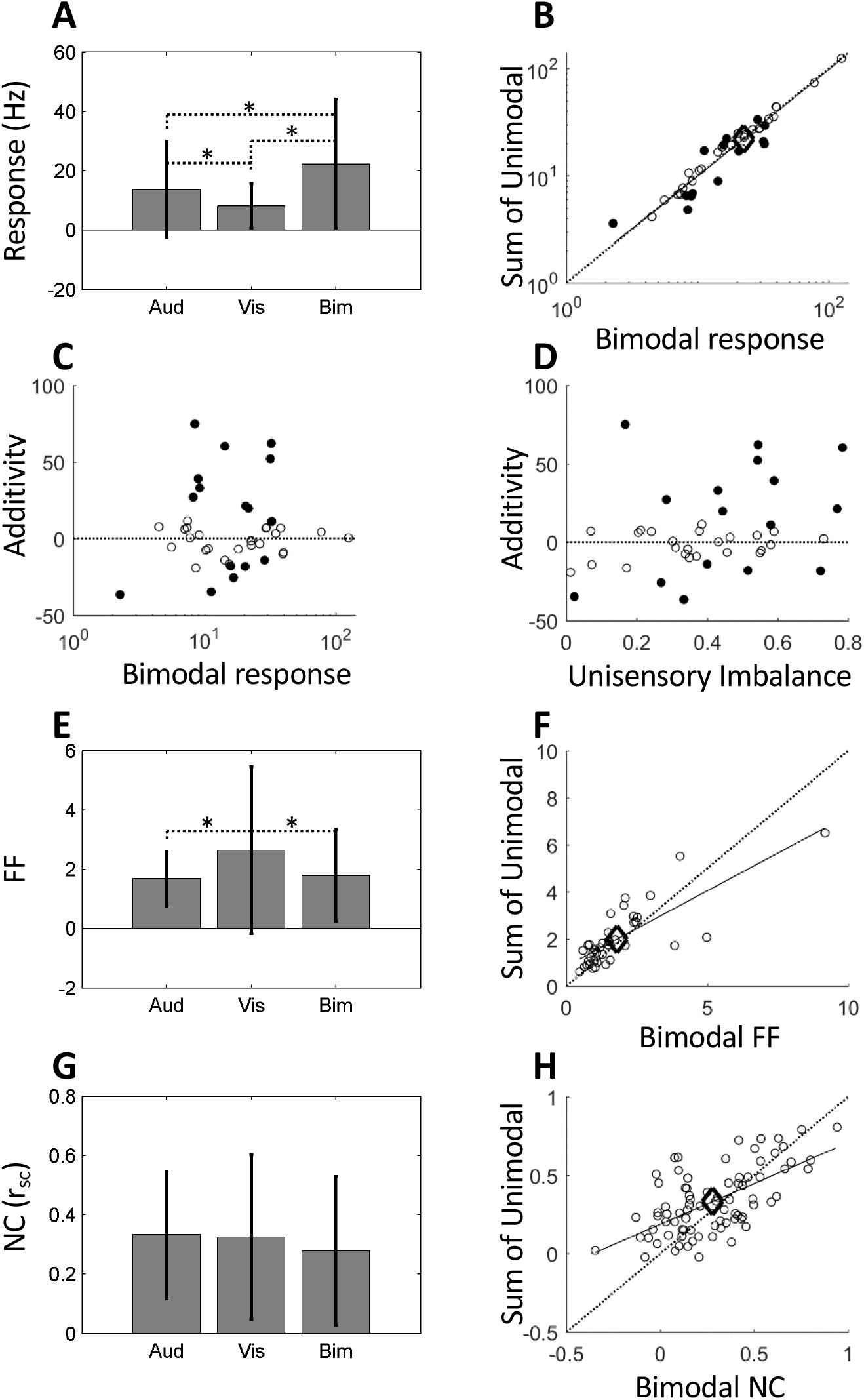
Bimodal integration. **A** Responses to auditory stimuli, measured as increases in firing rate above baseline, were significantly larger than responses to visual stimuli (13.8hz and 8.2hz respectively, n = 40, Wilcoxon signed rank, p < 0.05). Error bars denote one standard deviation, and asterisks denote significant differences between conditions. **B** Bimodal responses were, on average, highly correlated with and not significantly different from the sum of the unimodal components (sum = 0.98*bim + 0.11, rho = 0.98, means = 22.0 and 22.4 respectively, n = 40, Wilcoxon signed rank p > 0.05). Diamond indicates the means. Individual sites exhibiting significant non-additive responses are shown with filled circles (16 of 40). **C** Additivity was not significantly correlated with bimodal response magnitude (r = 0.06, p > 0.05). **D** Additivity was not significantly correlated with the absolute value of the unisensory imbalance (r = 0.31, p > 0.05). **E** FFs were significantly lower for auditory stimuli than for visual stimuli (1.68 and 2.64 respectively, n = 40, Wilcoxon sign rank test, p < 0.05). Bimodal FFs (mean = 1.79) were significantly lower than visual FFs (Wilcoxon sign rank test, p < 0.05) but not significantly different than auditory FFs. **F** Bimodal FFs were correlated with the FFs measured from the sum of the unimodal components, suggesting a large portion of the response variance occurs separately in each sensory stream (rho = 0.79, n = 40). While highly correlated, the bimodal FFs were also slightly lower than those from the sum of the unimodal components (1.79 and 1.99 respectively, p < 0.01). **G** Noise correlations were not significantly different for auditory, visual, or bimodal stimuli (0.33, 0.32, and 0.28 respectively, n = 80 ANOVA p > 0.05). **H** Bimodal NCs were correlated with NCs from the sum of the unimodal components (rho = 0.63, n = 80), and slightly lower for bimodal condition (0.28 and 0.33, t-test, p < 0.05).

To explore the response variance within each neuron we measured FFs while presenting visual, auditory and bimodal stimuli (Fig. 6E). Auditory responses exhibited lower FFs than visual responses (means of 1.68 and 2.64 respectively, p = 0.025 with a two-tailed signed rank test, n = 40), and the two were not significantly correlated. The bimodal FFs (mean of 1.79) were both significantly lower than visual FFs (p = 0.0002 with a two-tailed signed rank test) and correlated with them (Pearson’s r = 0.87, p < 0.0001). Conversely the bimodal FFs were not significantly different from, nor correlated with, the auditory FFs. To determine whether the bimodal FFs could be explained by linear integration of the unimodal components, the response magnitudes from each visual trial were added with an auditory trial and FFs were re-calculated. Auditory responses were trial-shuffled to calculate the FF with every combination of visual response, excluding the visual-auditory pair that occurred on the same trial number, thereby disrupting slow internal state variance that may be shared on flanking presentations. We found that even after shuffling to disrupt variance shared due to internal state fluctuations, bimodal FFs were still strongly correlated with FFs derived from linear integration of the auditory and visual signals (Fig. 6F, Pearson’s r = 0.79, p < 0.0001), while the bimodal FFs were slightly lower (means 1.79 and 1.99, p = 0.005 with a two-tailed signed rank test).

NCs values in this subset of 80 units used for bimodal analysis were similar in range (−0.35 to 0.94) and mean (0.28) to values reported for all pairs in Figure 5. There was not a significant effect of stimulus type on NC (Fig. 6G ANOVA p = 0.34, n = 80), but auditory and visual NCs were correlated with each other (Pearson’s r = 0.42, p = 0.0001) and with the bimodal values (auditory Pearson’s r = 0.52, visual r = 0.51). To determine whether additive interactions would explain the observed bimodal NCs, unimodal trials were summed and NCs calculated. Auditory responses for each cohort were trial-shuffled relative to visual responses, while maintaining trial alignment within the cohort so as not to affect within-modality NCs. Figure 6H shows that a strictly additive interaction would generate NCs that are moderately correlated with (Pearson’s r = 0.62, p < 0.0001), and slightly higher than (summed mean = 0.33, bimodal mean = 0.28, p = 0.036 with two-tailed T-test) the NCs evoked by bimodal stimuli. As shown in Figure 7, NC and FFs were correlated with each other for unimodal presentations (Aud: Pearson’s r = 0.41, p < 0.01; Vis: 0.34 p < 0.01) but uncorrelated during presentation of aligned bimodal stimuli (Pearson’s r = 0.21 p >0.05).

**Figure 7.**
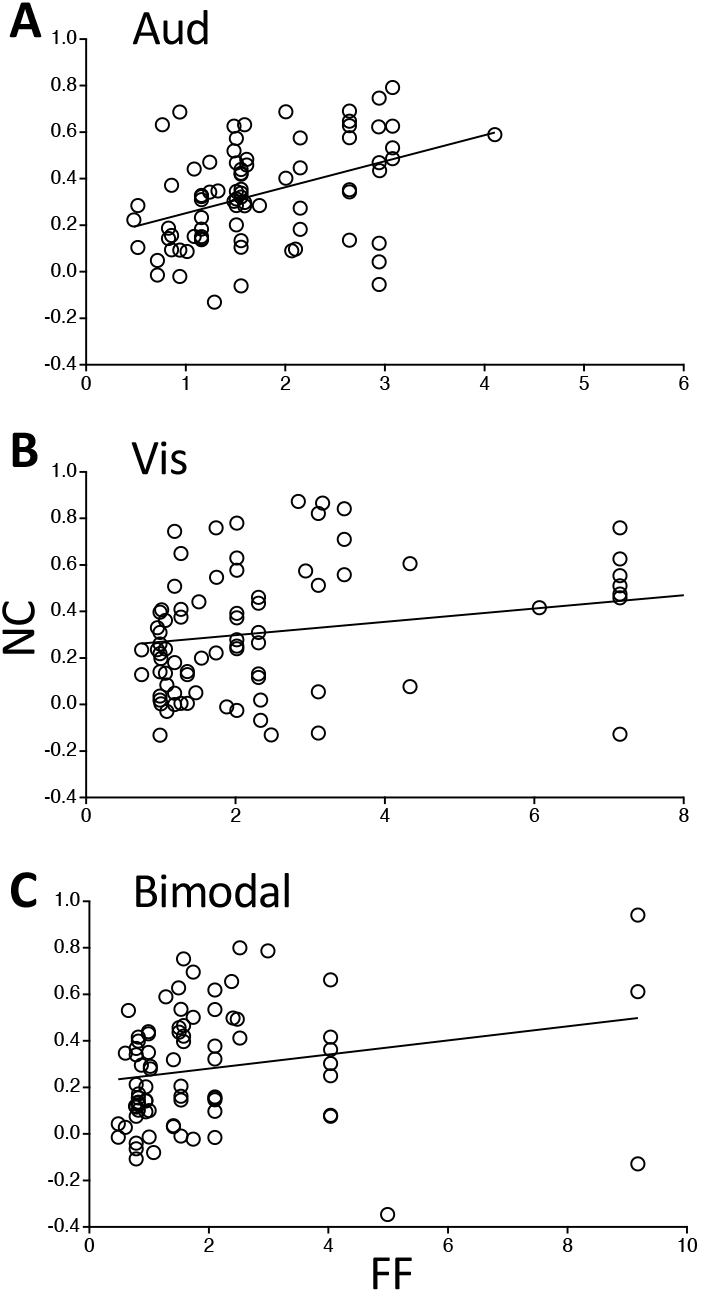
NC-FF correlations for aligned bimodal stimuli. **A** Auditory alone: n = 80, R^2^ = 0.17, p < 0.01; **B** Visual alone: n = 80, R^2^ = 0.12, p < 0.01; **C** Bimodal presentation: n = 80, R^2^ = 0.05, p > 0.05. p values determined using r to p conversion.

### Multisensory competition

To address how competing stimuli influence responses, FFs and NCs were recorded while presenting stimuli at the driver location in the overlapping portion of the cohort receptive fields, and at a location located 20° away, as shown in Figure 8A. We used three stimulus combinations: visual driver with visual competitor (unimodal), visual driver with auditory competitor (crossmodal), auditory driver with visual competitor (crossmodal). Auditory driver with auditory competitor was not used due to technical limitations of mimicking two sound sources with dichotic stimuli. The different sample sizes for different combinations are due to the fact that units were only included if they had a significant response to the driver and lacked a significant response to the competitor. Competing stimuli were effective at reducing responses to a driver stimulus regardless of the modality of the driver or the competitor (Fig. 8B, C, D) as has been previously shown (Mysore, Asadollahi, and Knudsen 2010). Interestingly, competitors decreased FFs for both auditory and visual drivers, but only when they were a different modality (Fig. 8F, G, K). Competitors of the same modality did not affect FFs (Fig. 8E, K). Likewise, NCs were significantly decreased for visual competitors when the driver was auditory (Fig. 8J, L), but significantly increased when the driver was visual (Fig. 8H, L). Consistent with these observations, there was a non-significant trend for auditory competitors to drive down NCs for visual drivers.

**Figure 8.**
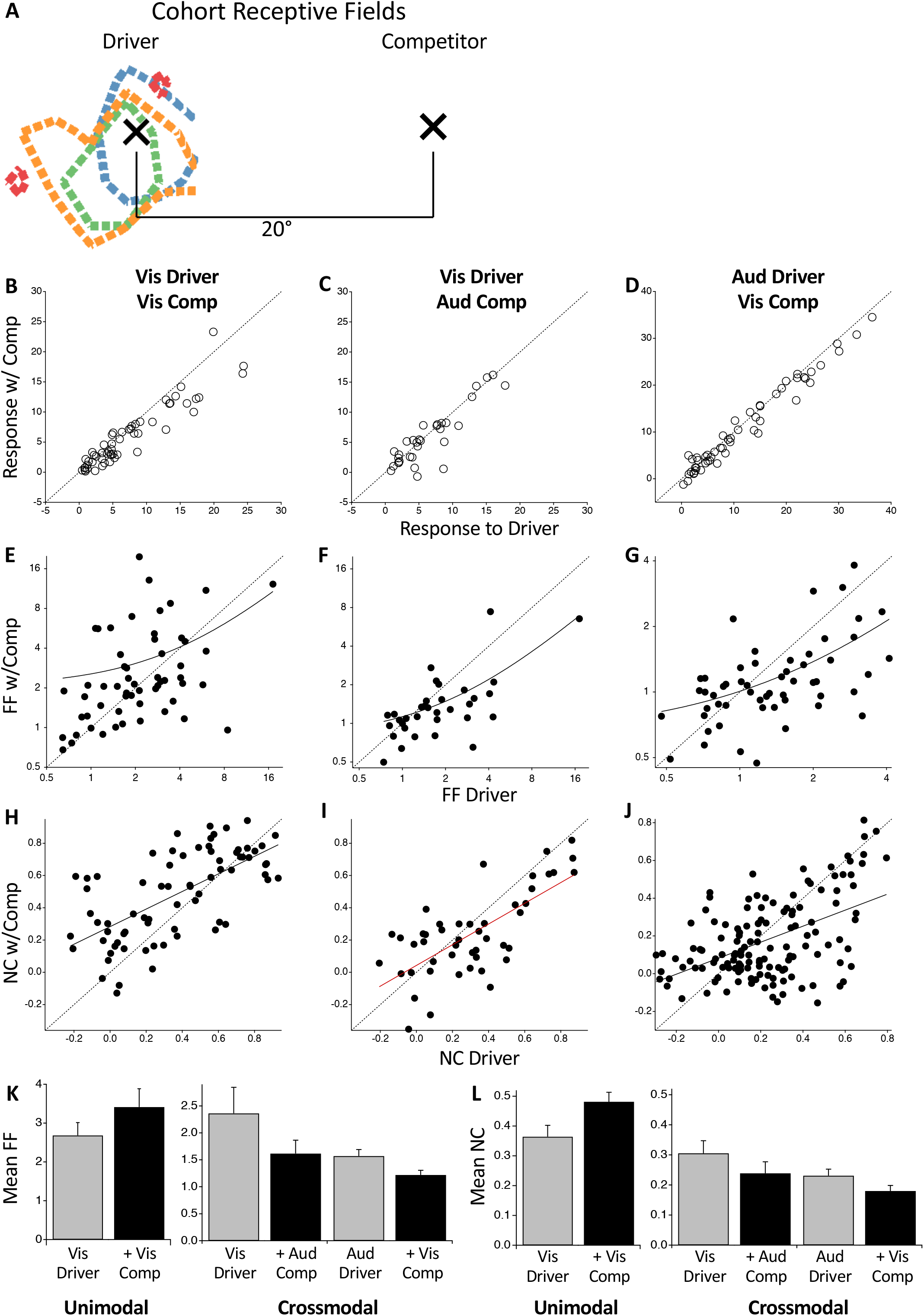
Effects of competing stimuli. **A** Auditory and visual stimuli were presented at driver locations and at competitor locations outside the spatial receptive field. Driver alone and competitor combinations were presented for 50 interleaved repetitions. **B, E, H** Visual driver with visual competitor (unimodal). **C, F, I** Visual driver with auditory competitor (crossmodal). **D, G, J** Auditory driver with visual competitor (crossmodal). Response correlations with and without the competitor: **B** n= 57, R^2^ = .93, p<0.01; **C** n = 34, R^2^ =.95, p<0.01; **D** n = 55, R^2^ =.99, p<0.01. FFs correlations with and without the competitor: **E** n= 57, R^2^ = .13, p<0.01; **F** n = 34, R^2^ =.49, p<0.01; **G** n = 55, R^2^ =.27, p<0.01. Linear regressions (solid lines) appear curved on log plots. NC correlations with and without the competitor: **H** n= 73, R^2^ = .43, p<0.01; **I** n = 47, R^2^ =.50, p<0.01; **J** n = 136, R^2^ =.25, p<0.01. Solid line is linear regression. **K** Vis-Vis competitor non-significantly increased FFs from 2.68 to 3.41 (n = 57, p > 0.05), whereas bimodal competitors significantly decreased FFs, from 2.36 to 1.62 for auditory competitors (n = 34, p < 0.05) and 1.57 to 1.22 for visual competitors (n = 55, p < 0.05). **L** Vis-vis competitor significantly increased NCs from 0.36 to 0.48 (n = 73, p < 0.05), whereas Vis driver + Aud competitor non-significantly decreased NCs, from 0.31 to 0.24 (n = 47, p > 0.05) and Aud driver + Vis competitor significantly decreased NCs from 0.23 to 0.18 (n = 136, p < 0.05). p values for correlations in **B-J** were determined using r to p conversion; p values in **K, L** were determined by Wilcoxon sign rank test.

For aligned stimuli, the correlations between NC and FF evident for unimodal presentations were disrupted by bimodal presentation (Fig. 7). In contrast, those correlations were not disrupted by non-aligned competitor stimuli regardless of competitor modality (Fig. 9).

**Figure 9.**
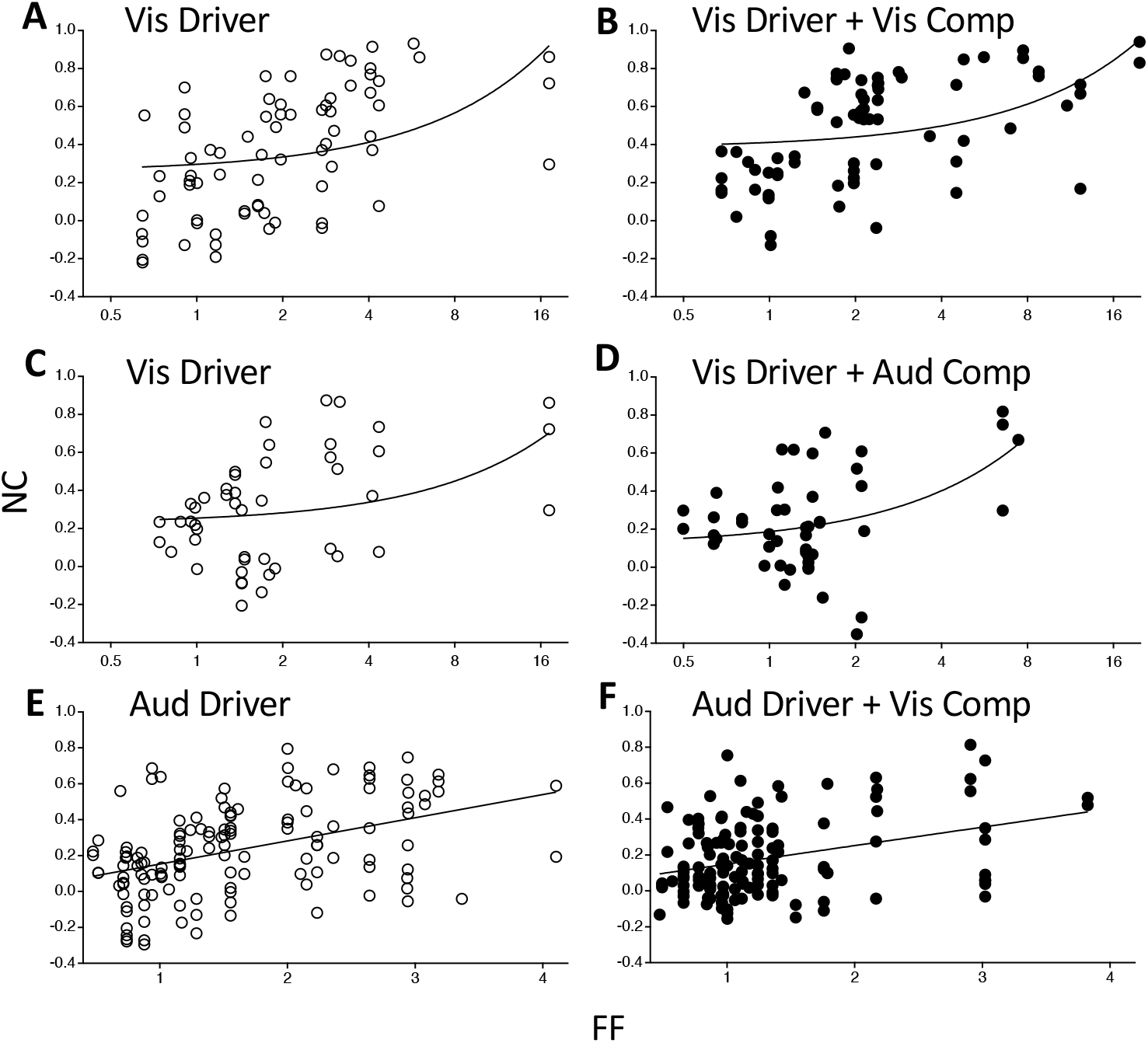
NC-FF correlations for non-aligned competitor stimuli. **A** Visual driver alone: n = 73, R^2^ .15, p < 0.01. **B** Visual driver with visual competitor: n = 73, R^2^ = .18, p < 0.01. **C** Visual driver alone: n = 47, R^2^ = .14, p < 0.01. **D** Visual driver with auditory competitor: n = 47, R^2^ = .20, p < 0.01. **E** Auditory driver alone: n = 136, R^2^ = .18, p < 0.01. **F** Auditory driver with visual competitor: n=136, R^2^ = .12, p < 0.01. The x-axis for **A-D** is log scale for clarity. p values determined using r to p conversion.

## DISCUSSION

The literature on multisensory processing in OTd is based largely on sequential recordings using single electrodes, which do not provide information on ensemble coding features - interneuronal signal and noise correlation. We addressed this gap using MEAs. The main findings are: (1) NC values are large and variable. (2) NCs decrease as tuning similarity (signal correlation) between neuronal pairs decrease, though the dependence is modest. (3) For spatially aligned stimuli, bimodal response magnitudes are well predicted from the sum of unimodal responses. Bimodal FFs and NCs can also be predicted by the additive integration of unimodal responses, although more variance is observed than for response magnitudes. (4) For spatially non-aligned stimuli, both unimodal and crossmodal competitors effectively drive down response magnitudes, consistent with previous (Mysore, Asadollahi, and Knudsen 2010, 2011) reports, but only crossmodal competitors could drive down FFs and NCs. In fact, unimodal competitors significantly increase NCs, and non-significantly increase FFs. These results indicate that the OT network is differentially wired to process unimodal vs crossmodal stimuli.

### NCs are large and variable

We examined noise correlations evoked by auditory (n = 182), visual (n = 181) or bimodal (n = 80) stimuli under conditions of light sedation. Evoked NC values ranged from -0.35 to 0.94, with a mean of 0.28, 0.25, and 0.28 for auditory, visual, and bimodal, respectively. These values are consistent with a prior report of 14 OTd pairs recorded in response to spontaneous activity (Netser, Dutta, and Gutfreund 2014). A recent study using tetrodes in OTd and auditory stimulation reported somewhat lower values, with mean NC values of 0.13 (Beckert, Pavao, and Pena 2017). We consider three possible explanations: (1) Our MEA cohorts contained neuronal pairs spanning a large range of tuning overlaps, whereas tetrode recordings largely reflected pairs of nearby neurons. However, both our data and Netser et al., found that NC values tend to decrease with increasing tuning difference, which should result in lower, not higher, mean values in the MEA data. (2) Differences in stimulation and analysis windows. Tetrode recordings employed 150ms stimulus and 300ms ISI, whereas MEAs used 250ms stimulus and 2500ms ISI. Given comparable firing rates, tetrode recordings are estimated to have produced ∼40% fewer spikes on average, which are expected to yield lower correlation values (Cohen and Kohn, 2011). Consistent with this, when a smaller analysis window was applied to the MEA data, NC values were slightly lower than when the entire acquisition window was interrogated (Figure 7). (3) Differences in anesthesia. Tetrode recordings were performed under ketamine anesthesia, MEA recordings under light sedation with nitrous oxide. While opioid anesthesia can produce artificially high NCs by inducing state changes (Ecker et al. 2014), we found no conclusive data relating NCs to ketamine or nitrous oxide.

The NC values we observed are at the high end of the range reported for cortical circuitry. A review of 26 studies of multiple areas of primate cortex revealed typical values of ∼0.1, with a maximum of 0.26 (Cohen and Kohn, 2011). The values reported here could reflect a large proportion of common drivers in the auditory and visual input streams, respectively. Unlike tonotopic maps in mouse auditory cortex in which neighboring neurons can exhibit starkly different frequency tunings despite global tonotopy (Rothschild and Mizrahi 2015; Kanold, Nelken, and Polley 2014), neighboring neurons in OT reliably exhibit similar tuning (Beckert, Pavao, and Pena 2017). This property is likely inherited from space-specific neurons (SSNs) in ICX via a point-to-point projection (Knudsen and Knudsen 1983). SSNs receive a broad range of ITD inputs from the adjoining lateral shell of the central nucleus of the inferior colliculus (ICCls) (Wagner, Takahashi, and Konishi 1987) yet synaptic connectivity is highest from map locations in ICCls encoding similar ITDs (DeBello and Knudsen 2001). This anatomical pattern would support integration of common drivers. Therefore, the observation of high NCs in OTd is consistent with the known functional and anatomical organization of the ICCls-ICX-OTd microcircuit.

Decoding the information represented by neuronal pairs using linear discriminant analysis demonstrated that low NCs (mean = 0.06) in owl forebrain (auditory arcopallium, AAR) limit the detrimental impact of NCs for pairs with positive signal correlations, resulting in better localization accuracy assuming a two channel rate-code scheme (Beckert, Pavao, and Pena 2017). This decoding scheme does not account for performance in OT, likely due to differences in the large-scale organization of ITD tuning between AAR and OT. Instead, a population decoder based on Bayesian inference accurately predicts head turn behavior from the average responses of OT neurons (Fischer and Pena 2011; Ferger et al. 2021). This decoder is also expected to be sensitive to signal and noise correlations. Our results regarding NCs and FFs from nearby and more distant pairs, and in response to aligned and non-aligned bimodal stimuli, provide new data to inform these population decoders.

### Rules for bimodal summation

Prior studies of multisensory integration in the avian OT or mammalian SC have demonstrated supralinear, linear or sublinear summation of auditory and visual inputs, depending on the position of the stimuli within their respective receptive fields (Ghose and Wallace 2014; Meredith and Stein 1996; Mysore, Asadollahi, and Knudsen 2010), unimodal response magnitudes (rule of inverse effectiveness) (Stanford, Quessy, and Stein 2005), relative magnitudes (sensory imbalance) and timing (Miller et al. 2015), and recent history of activation (Zahar, Reches, and Gutfreund 2009). The multisensory integration we observed was strikingly linear (Fig. 6). Comparison of the experimental conditions used across studies indicates that our results are broadly consistent with prior results.

An early study using ketamine anesthetized cats observed that most units (62/73) exhibited multisensory enhancement, defined as when the bimodal response exceed the sum of unimodal presentations (Meredith and Stein 1996). That effect, however, only occurred when both stimuli were positioned close to the center of their spatial receptive fields. In contrast, stimuli positioned near the receptive field borders showed no response enhancement while a single stimulus positioned outside the receptive field sometimes caused response suppression. In our dataset, SRFs of units in the cohort were often overlapping but rarely spot on, thus, individual stimuli were rarely optimal drivers. This regime is additive in both studies.

A related study found that when the unimodal response strengths were imbalanced and presented simultaneously, bimodal responses were additive (Miller et al. 2015). In contrast, if the stronger stimulus was presented first, or unimodal responses were well-balanced, summation tended to be supralinear. In our dataset, the only inclusion criterion was that both unimodal responses were significant above background, and therefore our population contained a wide spectrum of unimodal response magnitudes and relative magnitudes (i.e. imbalanced). This regime is additive in both studies. One difference is that none of the relatively few balanced units we recorded from exhibited supralinear behavior.

Two prior studies in owl OT revealed a similar range of integrative rules. Both used auditory and visual stimuli positioned at the center of the respective SRFs, and analyzed a mix of single and multiunits, both differences from our experimental design. One found supralinear summation but only when the auditory responses were weak (Mysore, Asadollahi, and Knudsen 2010). The other found linear to sublinear summation on average with a long supralinear tail (Zahar, Reches, and Gutfreund 2009). In summary, our data is broadly consistent with all previous studies in demonstrating that units can exhibit supralinear, linear or sublinear summation, and distinct in that our data does not reveal a clear predictive rule based on the relative magnitude and SRFs of unimodal responses.

### Crossmodal competitors drive down NCs while unimodal competitors do not

A major finding is that FFs and NCs decrease during most crossmodal competitions but do not for unimodal competitions, during which NCs in fact increase. It is tempting to argue that NC decreases during crossmodal interaction are due to the fact that auditory and visual signals arrive through separate pathways do not share bottom-up sensory correlations. However, if simply integrating across modalities decorrelated responses there should have been a significant decrease in NCs when presenting aligned auditory and visual stimuli, which was not observed (Fig. 6). Therefore, and given that neuronal pairs with positive signal correlation predominate the datasets (Fig. 5), these data on non-aligned competitors (Fig. 8) indicate that (a) the observed decrease in information-limiting NCs driven by crossmodal competition is predicted to improve localization accuracy for auditory targets with visual competitors, and (b) the observed increase in NCs driven by visual competitors on visual targets is predicted to harm localization accuracy. These predictions can be tested via computational studies using population decoding models.

Owls are known for their ability to hunt using sound alone (Payne, 1971), yet also possess high acuity night vision resulting from an all-rod area centralis (Fite, 1973; Fite and Rosenfield-Wessels, 1975). Crossmodal competitor stimuli encountered during natural hunting episodes could have selected, over evolutionary timescales, for optimal decoding in that condition.

## Acknowledgements

This work was supported by a NRSA Predoctoral Fellowship #F31 DC013708-02 to DJT; BRAIN-STIM grant from University of California Davis Office of Research and Behavioral Health Center for Excellence to WMD and National Institutes of Health Grants R01DC05640 (NIDCD) and R01NS104911 (BRAIN Initiative) to WMD.

